# Multivariate pattern analysis techniques for electroencephalography data to study interference effects

**DOI:** 10.1101/797415

**Authors:** David López-García, Alberto Sobrado, José M. G. Peñalver, Juan Manuel Górriz, María Ruz

## Abstract

A central challenge in cognitive neuroscience is to understand the neural mechanisms that underlie the capacity to control our behavior according to internal goals. Flanker tasks, which require responding to stimuli surrounded by distracters that trigger incompatible action tendencies, are frequently used to measure this conflict. Even though the interference generated in these situations has been broadly studied, multivariate analysis techniques can shed new light into the underlying neural mechanisms. The current study is an initial approximation to adapt an interference Flanker paradigm embedded in a Demand-Selection Task to a format that allows measuring concurrent high-density electroencephalography. We used multivariate pattern analysis (MVPA) to decode conflict-related neural processes associated with congruent or incongruent target events in a time-frequency resolved way. Our results replicate findings obtained with other analysis approaches and offer new information regarding the dynamics of the underlying mechanisms, which show signs of reinstantiation. Our findings, some of which could not had been obtained with classic analytical strategies, open novel avenues of research.

## 1 Introduction

Cognitive control comprises a set of mechanisms that allow humans to behave according to their internal goals while ignoring distracting information.^1^ The flanker task, where participants respond to the direction of an arrow surrounded by other distracting arrows, is among the most used in the field. The main result of this task is the so-called interference or conflict effect, where responses are slower and less accurate in incongruent (when the direction of the distracters is opposite to the target) vs. congruent trials. In the current study, we employed a task that measured interference effects in the context of effort avoidance.^2^ Cognitive control involves effort, which is costly and partly aversive, and thus humans usually avoid it if given the chance. In Demand-Selection Tasks (DST),^2^ participants tend to choose the easy option over the hard one. The tendency to avoid the hard option seems partly due to the cost of overcoming the increased cognitive control required when responding to incongruent situations. However, the neural underpinnings of this effect are not well understood.

The majority of Electroencephalography (EEG) studies of the interference effect have analyzed Event-Related Potentials (ERPs), focusing on the N2 component. Besides, studies employing frequency analyses have shown Theta and Delta band involvement. Other authors^3^ have proposed a link between the ERPs and modulations in the Delta-Theta band of frequency.

Supervised Machine Learning algorithms, more specifically Linear Support Vector Machines (LSVM)(Vapnik, 1979) ^4,5^ have been widely applied in clinical settings such as computer-aided diagnosis of Alzheimer’s disease,^6–9^ automatic sleep stages classification^10,11^ or automatic detection of sleep disorders.^12^ In recent years, these Machine Learning-based analyses, in conjunction with neuroimaging techniques such as functional Magnetic Resonance Imaging (fMRI), Electroencephalography or Magnetoencephalography (MEG), have gained popularity in Cognitive Neuroscience.^13–18^

One of the most remarkable advantages of Multivariate Pattern Analysis (MVPA) versus univariate approaches is its sensitivity in detecting subtle changes in the patterns of neural activity.^19^ When applied to fMRI data, the poor temporal resolution of the signal prevents an accurate study of how cognitive processes unfold in time. In contrast, when applied to M/EEEG signals, MVPA have been useful to uncover the neural dynamics of face detection,^20^ the process of memory retrieval,^21^ the representational dynamics of task and object processing in humans^22^ or the representation of spoken words in bilingual listeners.^23^ In the same line, time-resolved MVPA presents an opportunity to categorize the temporal sequence of the neural processes underlying the interference effect. Furthermore, the relationship between these and the Theta frequency modulations can be better understood using this approach. The goal of the current study is to present a set of methodological MVPA tools that allow to study and decode the conflict-related neural processes underlying interference effects, in a time-frequency resolved way.

## 2 Materials and methods

### 2.1 Paradigm and data acquisition

#### 2.1.1 Participants

Thirty-two healthy individuals (21 females, 29 right-handed, mean age = 24.65, SD = 4.57) were recruited for the experiment. Participants had normal or corrected-to-normal vision and no neurological or psychiatric disorders. All of them provided informed, written consent before the beginning of the experiment and received a 10-euro payment or course credits in exchange for their participation. The experiment was approved by the Ethics Committee of the University of Granada.

#### 2.1.2 Stimuli and apparatus

Stimuli presentation and behavioral data collection were carried out using MATLAB (MathWorks) in conjunction with Phychtoolbox-3.^24^ The visual stimuli were presented in an LCD screen (Benq, 1920×1080 resolution, 60 Hz refresh rate) and placed 68.31*±*5.37 cm away of participant’s Glabella, in a magnetically shielded room. Using a photodetector, the stimuli onset lag was measured at 8ms, which corresponds to half of the refresh rate of the monitor. Triggers were sent from the presentation computer to the EEG recording system through an 8-bit parallel port and using a custom MATLAB function in conjunction with inpoutx64 driver,^25^ a C++ extension (mex-file) that uses native methods to access low-level hardware in MATLAB (I/O parallel ports).

Cues consisted of two squares of two different colors (red/green and yellow/blue, in different blocks) stacked and presented at the center of the screen (visual angle ∼5 degree). In forced blocks, a small white indicator (circle 50% or square 50%) appeared on top of the color that had to be chosen. In voluntary blocks, this indicator appeared between the two colored squares (see Figure 1). Each target stimulus consisted of five arrows pointing left or rightwards, which were displayed at the center of the screen (visual angle ∼6 degree). The color of the target stimulus was the same as the cue previously selected.

**Figure 1:**
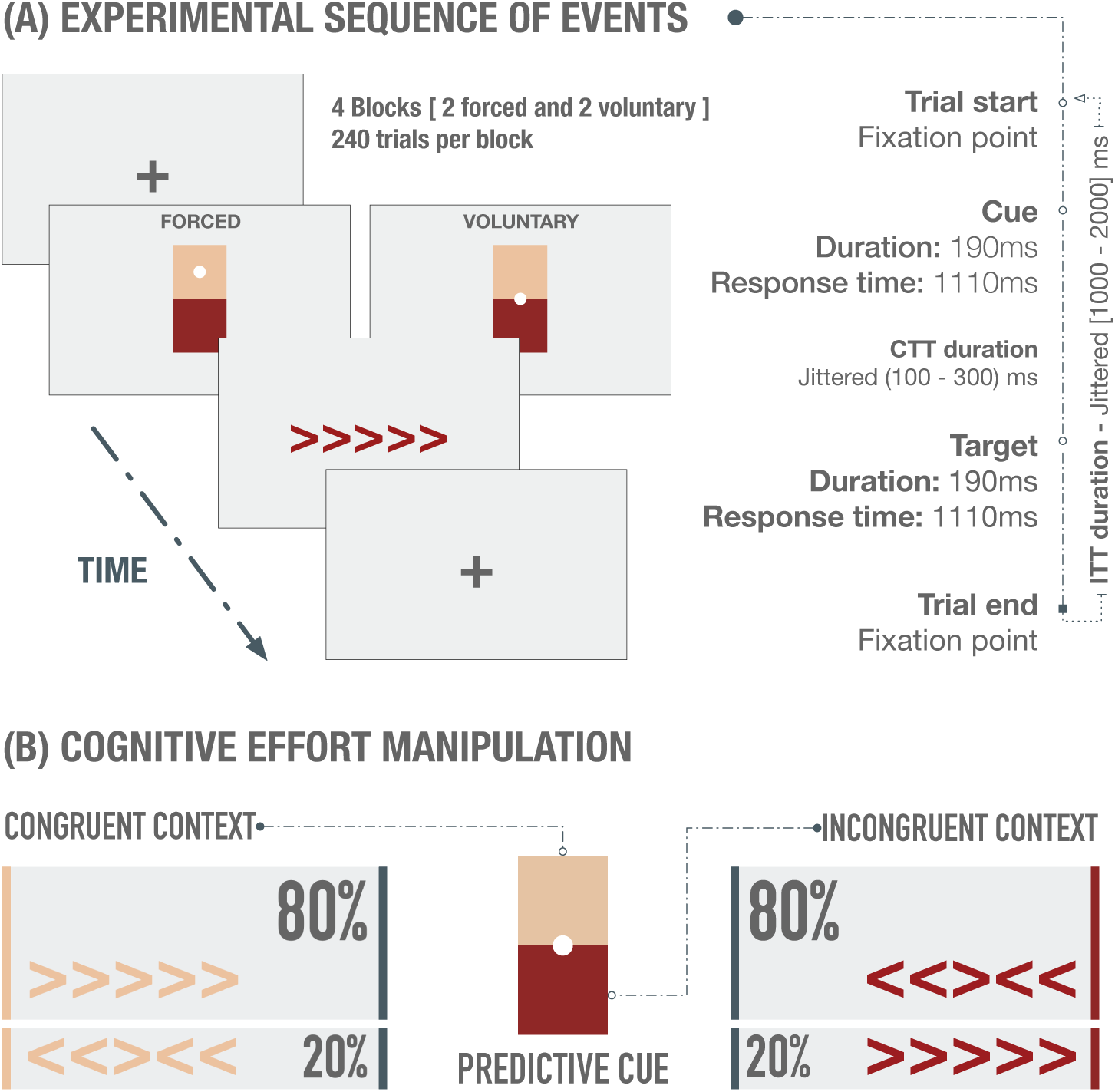
**(A)** Experimental sequence of events in case of a correct response on both cue and flanker stimuli. Each trial started with a fixation point, followed by a cue, which acted as a selector of the difficulty of the upcoming Flanker target. Participants had to choose (freely or forced, depending on the block type) the posible color of the upcoming target stimulus, which was associated with either high (difficult) or low (easy) probability of incongruent trials. Finally, after a variable time interval (100-300ms) the target stimulus appeared and participants had to respond to the orientation of the central arrow. Another variable time interval appeared before the beginning of the next trial. The cue and the target stimuli remained on screen for 190ms. **(B)** Cognitive effort was manipulated through the percentage of congruent and incongruent trials. Each cue color was associated with the high or low conflict contexts.

#### 2.1.3 Procedure

The Color-Based Demand-Selection Task (DST) (Figure 1 a), modified from,^2^ consisted of a cue-target sequence arranged in four blocks (2 forced and 2 voluntary). In voluntary blocks, participants were required to freely choose one of the two colors available, which indicated the difficulty of the upcoming task. In forced blocks, a small white indicator appeared on top of the color that had to be chosen. The color of the target stimulus was the same as the cue previously selected and participants were required to discriminate the orientation (right or left) of a central arrow target surrounded by arrows pointing at the same (compatible distractors) or opposite (incompatible distractors) directions.

Our task was built following a 3-way factorial design, containing the following within-subjects independent variables: (1) Stimulus type (congruent/incongruent); (2) Block type (forced/voluntary) and (3) Context (easy/difficult). The task difficulty manipulation was based on the proportion of congruent and incongruent trials, with the easy contexts presenting 80% of congruent and 20% of incongruent trials, and the difficult task context the opposite proportion. Within forced blocks, half of the trials corresponded to the easy context and the remaining to the difficult one (maintaining, within each condition, the proportion of congruent and incongruent trials). On voluntary blocks, however, participants freely chose the context and no experimental control could be exerted upon this variable.

Participants were instructed to respond as fast and accurately as possible, and to not choose color based on personal preference. They were unaware of the cognitive effort manipulation. To preserve the signals as clean as possible and remove the least number of trials, participants were encouraged to remain as still and relaxed as possible, avoiding face muscle activity and eye movements, but blinking normally. The order of the blocks, cue colors, response keys and color-conflict context mappings were counterbalanced across participants. There were 4 blocks, 240 trials per block, and the total recording session lasted ∼ 90min. Before the experimental session, participants performed a brief practice to familiarize themselves with the task (4 blocks, 20 trials per block, practice duration ∼20min).

#### 2.1.4 EEG acquisition and preprocessing

High-density electroencephalography was recorded from 65 electrodes mounted on an elastic cap (actiCap slim, Brain Products) at the Mind, Brain, and Behavior Research Center (CIMCYC, University of Granada, Spain). The TP9 and TP10 electrodes were used to record the electrooculogram (EOG) and were placed below and next to the left eye of the participant. Impedances were kept below 5*k*Ω. EEG activity was referenced online to the FCz electrode and signals were digitized at a sampling rate of 1KHz.

Electroencephalography recordings were average referenced, downsampled to 256Hz, and digitally filtered using a low-pass FIR filter with a cutoff frequency of 120Hz, preserving the phase information. The recording amplifiers have an intrinsic lower cutoff frequency of 0.016Hz (time constant *τ* = 10s).

No channel was interpolated for any participant. EEG recordings were epoched [-1000, 2000ms centered at onset of the target arrows] and baseline corrected [-200, 0ms], and data was extracted only from correct trials. To remove blinks from the remaining data, Independent Component Analysis (ICA) was computed using the *runica* algorithm in EEGLAB,^26^ excluding TP9 and TP10 channels. Artifactual components were rejected by visual inspection of raw activity of each component, scalp maps and power spectrum. Then, an automatic trial rejection process was performed, pruning the data from non-stereotypical artifacts. The trial rejection procedure was based on (1) abnormal spectra: the spectrum should not deviate from baseline by *±*50dB in the 0-2 Hz frequency window, which is optimal for localizing any remaining eye movements, and should not deviate by -100dB or +25dB in 20-40Hz, useful for detecting muscle activity (∼1% of the total sample was rejected); (2) improbable data: the probability of occurrence of each trial was computed by determining the probability distribution of values across the data epochs. Trials were thresholded, in terms of *±*6*SD*, and automatically rejected (∼6% of the total sample); (3) extreme values: all trials with amplitudes in any electrode out of a *±*150*µ*V range were automatically rejected (∼3% of the total sample).

#### 2.1.5 Final dataset description

The final dataset for our binary classification problem is shown in Table 1, where *N* is the initial number of trials per participant and class, *N*_*r*_ represents the number of remaining correct trials after the trial rejection stage and *N*_*b*_ is the final number of trials used for classification per participant (after downsampling the majority class in order to get balanced datasets).

**Table 1:** Number of observations of the final dataset

| Observations per participant | $N$ | $N_r$ | $N_b$ |
| --- | --- | --- | --- |
| Congruent class | 480 | $426 \pm 49$ | $359 \pm 52$ |
| Incongruent class | 480 | $368 \pm 59$ | $359 \pm 52$ |
| Total number of observations | $N_{Tr}$ | | $N_{Tb}$ |
| Congruent class | 13644 |  | 11505 |
| Incongruent class | 11782 |  | 11505 |

#### 2.1.6 Behavioral data analysis

Reaction time (RT) and error rates were registered for each participant. Before the statistical analysis, the first trial of each block, trials with choice errors and trials after errors were filtered out.^27^ Finally, RT outliers were also rejected using a *±*2.5 SD threshold, calculated individually per participant and condition. To analyze behavioral data (accuracy and reaction times) we conducted a repeated-measures ANOVA in IBM SPSS Statistics Software (v.20). Post hoc tests were carried out on the significant interactions using a Bonferroni correction for multiple comparisons.

### 2.2 Multivariate pattern analysis

The MVPA for the decoding analysis was performed in MATLAB by a custom-developed set of linear Support Vector Machines, trained to discriminate between congruent and incongruent target stimuli. To avoid skewed classification results, the datasets were strictly balanced, by downsampling the majority class to match the size of the minority one. In addition, class size was set as a factor of *k*, the total number of folds in the cross-validation stage. Accordingly, each fold was composed by exactly the same number of observations, avoiding any kind of bias in the results. The rest of the classification parameters remained by default.

**Figure 2:**
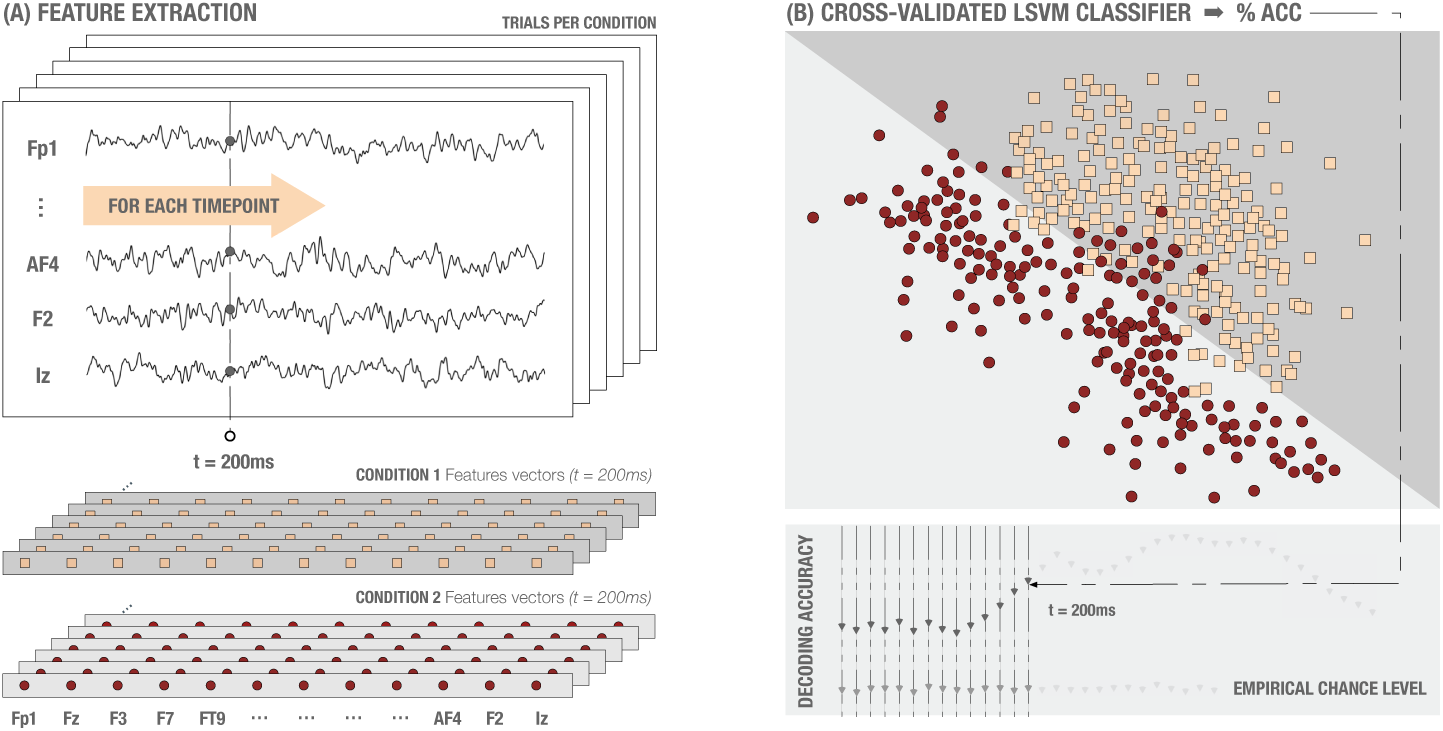
**(A)** Feature extraction process in simulated data. The feature vectors of each condition and time point consisted of a z-scored voltage array for all the scalp electrodes. For an improved SNR, several trials were averaged before feature extraction. **(B)** Cross-validated LSVM classifier. For each time point, an LSVM was trained and tested (stratified k-fold cross-validation, k=5). Chance level was calculated by permuting the labels.

#### 2.2.1 Feature extraction

To obtain the classification performance in a time-resolved way, the feature vectors were extracted as shown in Figure 2. The classification procedure, for each participant, ran as follows: (1) For each timepoint and trial, we generated two feature vectors (one for each condition or class) consisting of the raw potential measured in all electrodes (excluding EOG electrodes: TP9 and TP10). (2) Features vector containing raw potential values were normalized (z-score).

#### 2.2.2 Supertrial generation

Due to the noisy nature of the EEG signal, a trial averaging approach was carried out during the feature extraction stage. This approach increases the signal-to-noise ratio (SNR),^28^ improves the overall decoding performance and also reduces the computational load. Each participant’s dataset was reduced by randomly averaging a number of trials *t*_*a*_ belonging to the same condition. The value of *t*_*a*_ is a trade-off between an increased classification performance (due to an increased SNR) and the variance in the classifier performance, since reducing number of trials per condition typically increases the variance in (within-participant) classifier performance.^29^ Therefore, the optimal number of trials to average depends on the dataset, taking into account that averaging more trials does not increment the decoding performance linearly.

#### 2.2.3 Feature selection

As mentioned in section 2.2.1, **X**_n×p_ datasets are generated for each participant and timepoint, where n is the number of trials (observations) and p the total number of electrodes (variables or features). In machine learning, feature selection techniques, also known as dimension reduction, are a common practice to reduce the number of variables in high-dimensional datasets (Figure 3). Principal Component Analysis (PCA) is probably the most used and popular multivariate statistical technique and it is used in almost all scientific disciplines,^30^ including Neuroscience.^31^

PCA is a linear transformation of the original dataset in an orthogonal co-ordinate system in which axis coordinates (principal components) correspond to the directions of highest variance sorted by importance. To compute this transformation,^32^ each row vector **x**_*i*_ of the original dataset **X** is mapped to a new vector of principal components **t**_*i*_ = (*t*_1_, …, *t*_*l*_), also called *scores*, using a p-dimensional *coefficient* vector **w**_*j*_ = (*w*_1_, …, *w*_*p*_). For dimension reduction, *l < p*.

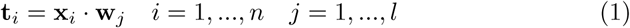

To maintain the model’s performance as fair as possible, in our study PCA was computed only for training sets **X**_training_, independently for each fold inside the cross-validation procedure. Once PCA for the corresponding training set was computed and the model was trained, the exact same transformation was applied to the test set **X**_test_ (including centering, *µ*_*training*_). In other words, the test set was projected onto the reduced feature space obtained during the training stage. According to equation (1), this projection is computed as follows:

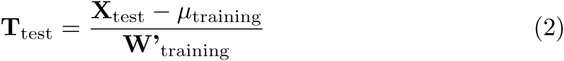

Feature selection techniques such PCA usually imply an intrinsic loss of spatial information, e.g. data projected from the sensor space onto the reduced PCA features space. Therefore, PCA presents a trade-off between dimension reduction and results’ interpretation. If PCA is computed, the spatial information of each electrode is lost, which means that, for example, we cannot directly analyze which electrodes are contributing more to the decoding performance.

#### 2.2.4 Model’s performance evaluation

To evaluate classification models in neuroscience, performance is usually measured employing mean accuracy.^33^ However, mean accuracy may generate systematic biases in situations with very skewed sample distributions and overfitting one single class should be avoided. Therefore, nonparametric and criterion-free estimates, such as the Area Under the ROC Curve (AUC) have been proved as a better measure of generalization in these situations.^34^ The AUC provides a way to evaluate the performance of a classification model. The larger the area, the more accurate the classification model is, and it is computed as follows:

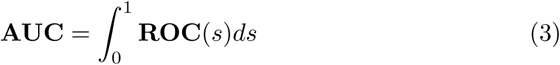

The ROC curve is one of the most important evaluation criteria, which shows how capable the model is in distinguishing between conditions, by facing the sensitivity (*True Positive Rate*, TPR) against 1-specificity (*False Positive Rate*, FPR). In this study, we employed both accuracy methods, to replicate a common approach in literature, and ROC curves and AUC, to provide a more informative measure.

To evaluate the performance of our model, LSVMs were trained and validated, resulting in a single performance value for each timepoint and participant. The classification performance at the group level was calculated by averaging these values across participants. The chance level was calculated following the former analysis but using randomly permuted labels for each trial.

The generalization ability of our model was estimated through a Cross-Validation (CV) approach (stratified k-fold, *k = 5*), which is a well-established and a widely implemented technique to preserve complex models from overfitting.

Moreover, some important aspects are worth being highlighted. The use of CV approaches often leads (particularly in Neuroscience) to small sample sizes and a high level of heterogeneity when conditions are splitted into each fold, causing among other things a high level of classification variability. To address these problems, a recent study^35^ considered the use of the resubstitution error estimate when using LSVM (in small sample sizes and low dimensional scenarios), proposing a novel analytic expression for the upper bound on the actual risk *γ*_*emp*_(*l, d*) for a range of sample sizes *l*, dimensions *d* and any significance *η <* 0.05 (Figure 8). Therefore, the difference between the actual error and the resubstitution error is bounded by the actual risk *γ*_*emp*_, which is computed as follows:

**Figure 3:**
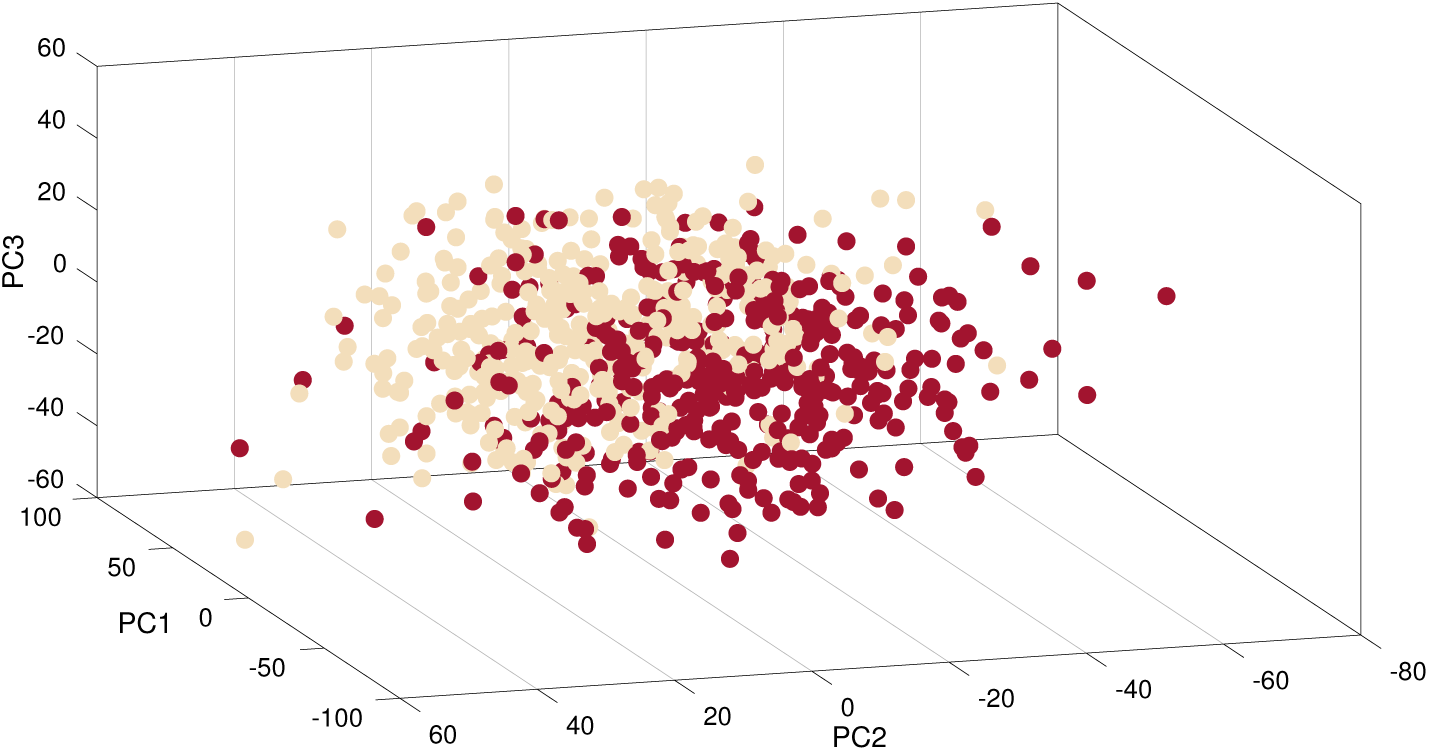
Dimension reduction in real datasets. 3D representation of the three first PCA components for congruent vs. incongruent trials [example participant, t = 421ms after Flanker stimulus onset]

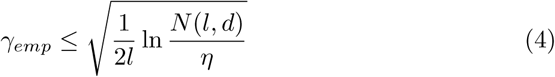

where N is defined in^35^ as:

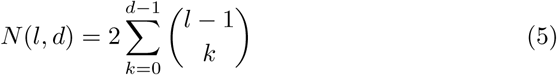

Resubstitution has been proved competitive in some heterogeneous-data scenarios with CV approaches not only in terms of accuracy but also in computational load.^36^ The proposed solution has been recently applied in clinical settings studying autistic patterns^37^ or Alzheimers Disease.^38^ The scenario previously mentioned (linear classifiers, small sample size and low dimensional space) seems to fit perfectly with our study setup, therefore, the use of the resubstitution error estimate in Cognitive Neuroscience studies is worthy of consideration.

#### 2.2.5 Optimization of SVM hyperparameters

A search-grid based optimization of the misclassification cost parameter *C* was carried out using five-fold cross-validation on the training set:

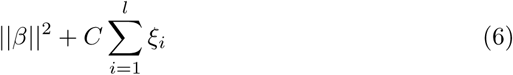

where *C* is a constant which modulates the trade-off between the training error and the complexity of the model and the vector *β* contains the coefficients that define an orthogonal vector to the hyperplane.

### 2.3 Temporal generalization matrix

Temporal generalization analyses are used to evaluate the stability of the brain patterns along time, by training the model in one temporal point and testing its capability to correctly discriminate between conditions in the remaining temporal window. This process is repeated for every timepoint. In our study, classification performance was assessed through a cross-validation technique (stratified k-fold cross-validation, k=5). For each timepoint, the classifier was trained with **X**_training_ dataset and tested with **X**_test_ in the remaining points of the temporal window. This process was repeated k times obtaining the final decoding accuracy.

An above-chance discrimination rate outside the diagonal of the matrix suggests that the same activity pattern is sustained in time. However, if there is no evidence of temporal generalization, separate patterns of activity can be assumed.^22^

### 2.4 Multivariate cross-classification

The ability of MVPA to detect subtle differences in brain activity patterns can be used to study how these patterns are similar across different cognitive contexts. In other words, the consistency of the information across different sets of data can be analyzed. To this end, classification algorithms are trained with one set of data and the consistency is assessed by testing the model with another dataset, belonging to a different experimental condition. This technique is called *Multivariate Cross-Clasification*^19^ (MVCC) and is growing in popularity in recent years.^39–41^

The fact that the training and test sets are different eliminates the need to use cross-validation techniques. However, several considerations have to be taken into account. First, the right choice of classification direction, that is, which set is used for training and which one for testing. The result of the classification could differ if, for instance, the signal-to-noise ratio is quite different across datasets, that is to say, differences in classes separability across datasets and an asymmetry in the generalization direction.^42^ For this reason, reporting results in both directions is highly recommended.

In this study, MVCC was used to analyze if the neural patterns associated with the congruency effect are similar across voluntary and forced blocks. For that, classifiers were trained with data of forced blocks and tested in voluntary blocks, and *vice versa*. In addition, a temporal generalization matrix was also computed to study the similarity across block types and time. Feature selection in MVCC analysis also requires some additional considerations, as features selected for the training set could not be the optimal selection for the test set. To avoid possible skewed results, no feature selection was computed in our study.

**Figure 4:**
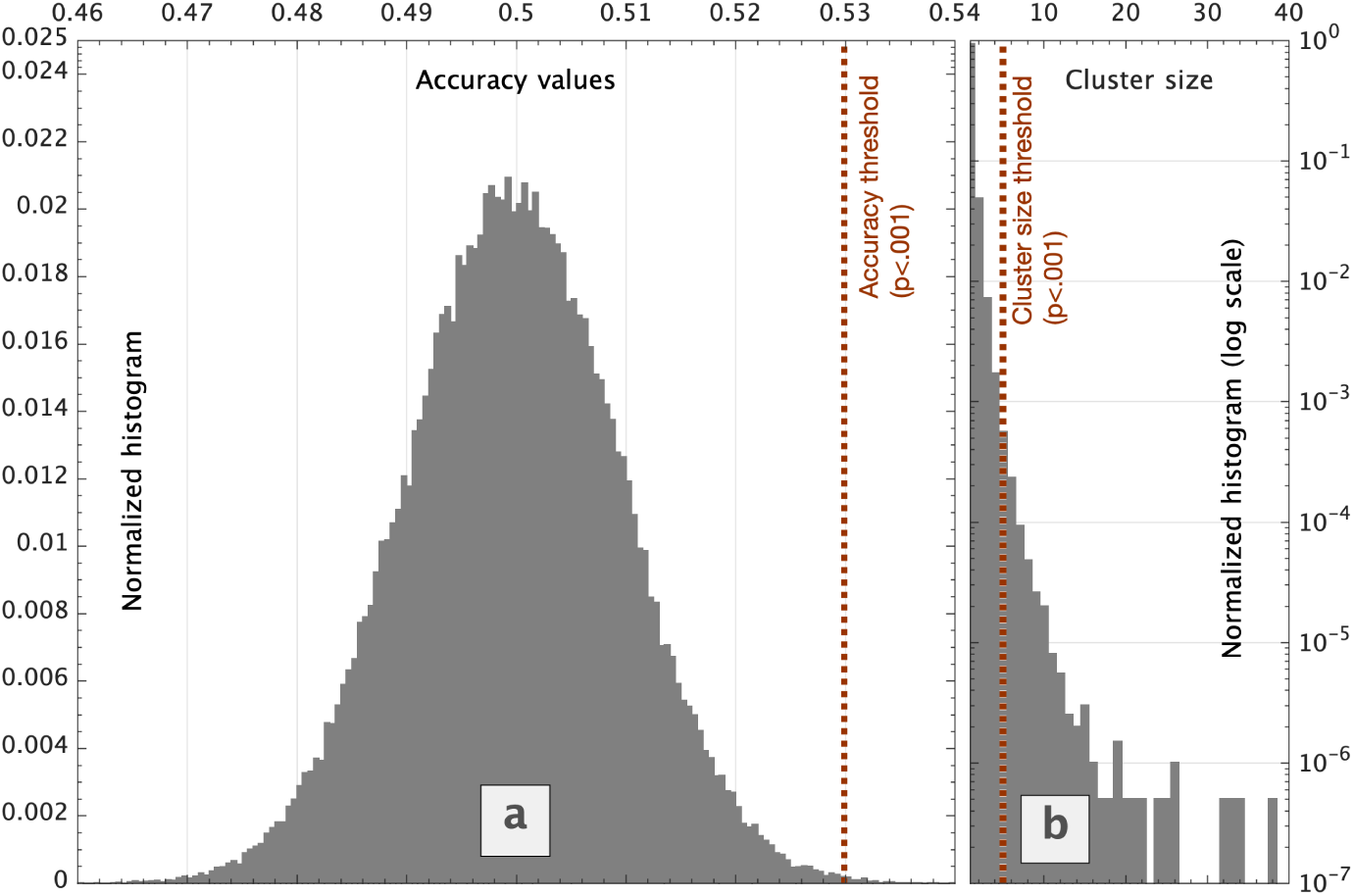
Accuracy **(a)** and cluster size **(b)** null distributions. The vertical dotted line represents the threshold corresponding to a very low probability to obtain significant results by chance. This threshold correspond to a p-value below 0.001 for both distributions.

### 2.5 Statistical analysis

Applying t-test statistics on multivariate results is an unsuitable approach to draw statistical inferences at the group level.^43^ For that reason, the use of cluster-based non-parametric permutation methods is widespread, not only in fMRI^44–47^ but more recently also in M/EEG studies.^48–51^ In our study, a nonparametric cluster-based permutation approach, proposed in^43^ for fMRI data, was adapted and implemented for the statistical analysis.

We thresholded the decoding accuracy obtained with an empirical accuracy null distribution, calculated by means of a combined permutation and bootstrapping technique. First, at the single-subject level, 100 randomly permuted accuracy maps were generated. To draw statistical inferences at the group level, we randomly drew one of the previously calculated accuracy maps for each participant. This selection was group-averaged and the procedure was repeated 10^5^ times, generating 10^5^ permuted group accuracy maps.

Next, for each timepoint we estimated the chance distribution of accuracy values and determined the accuracy threshold (99.9th percentile of the right-tailed area of the distribution), which corresponds to a very low probability to obtain significant results by chance.

Then, we searched and collected clusters of timepoints exceeding the previously calculated threshold in all the 10^5^ permuted accuracy maps, generating the normalized null distribution of cluster sizes. Finally, we applied a correction for multiple comparisons (FDR) to obtain the smallest cluster size to be considered significant.

### 2.6 Frequency contribution analysis

The contribution of each frequency band to the overall decoding performance was assessed through an exploratory sliding filter approach. We designed a band-stop FIR filter using `pop firws` EEGLab function (2Hz bandwidth, 0.2Hz transition band, 2816 filter order, Blackman window) and pre-filtered the EEG data (120 overlapped frequency bands, between 0-120Hz and linearly-spaced steps) producing 120 filtered versions of the original EEG dataset. The former time-resolved decoding analysis (congruent vs. incongruent, ta = 10) was repeated for each filtered version and the importance of each filtered-out band was quantified computing the difference maps in decoding performance between the filtered and the original decoding results. Significant clusters were found applying the proposed cluster-based permutation test to filtered-out datasets, generating accuracy null distributions for each time-frequency point.

With the purpose of obtaining better frequency resolution in lower bands, the previous analysis was repeated for frequencies between 0-40Hz in 120 overlapped and logarithmically spaced steps.

## 3 Results and discussion

### 3.1 Behavioral results

The behavioral results of reaction times replicate well-known conflict effects linked to context-dependent congruency,^2,27^ with a significant interaction of Context × Stimulus Type 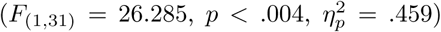. Planned comparisons showed significant differences between congruent and incongruent trials for both the easy 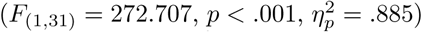 and the difficult contexts 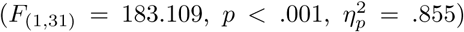 with larger differences in reaction times in the easy (congruent trials: *M* = 0.465, *SD* = 0.13; incongruent trials: *M* = 0.560, *SD* = 0.15), compared to the difficult context (congruent trials: *M* = 0.474, *SD* = 0.13; incongruent trials: *M* = 0.553, *SD* = 0.14).

### 3.2 Electrophysiological results

The electrophysiological analyses (Figure 5a) show significant differences (p*<*.001, cluster corrected) in activity patterns for congruent vs. incongruent trials, peaking at 375ms after the stimulus onset. At this point, the classifier accurately predicted (*>* 80%) if participants were responding to congruent or incongruent trials. Table 2 reports the variations in classification performance for averages of different number of trials. The SVM hyper-parameter C was optimized, slightly increasing the decoding performance, however, the computation time required increased significantly.

**Figure 5:**
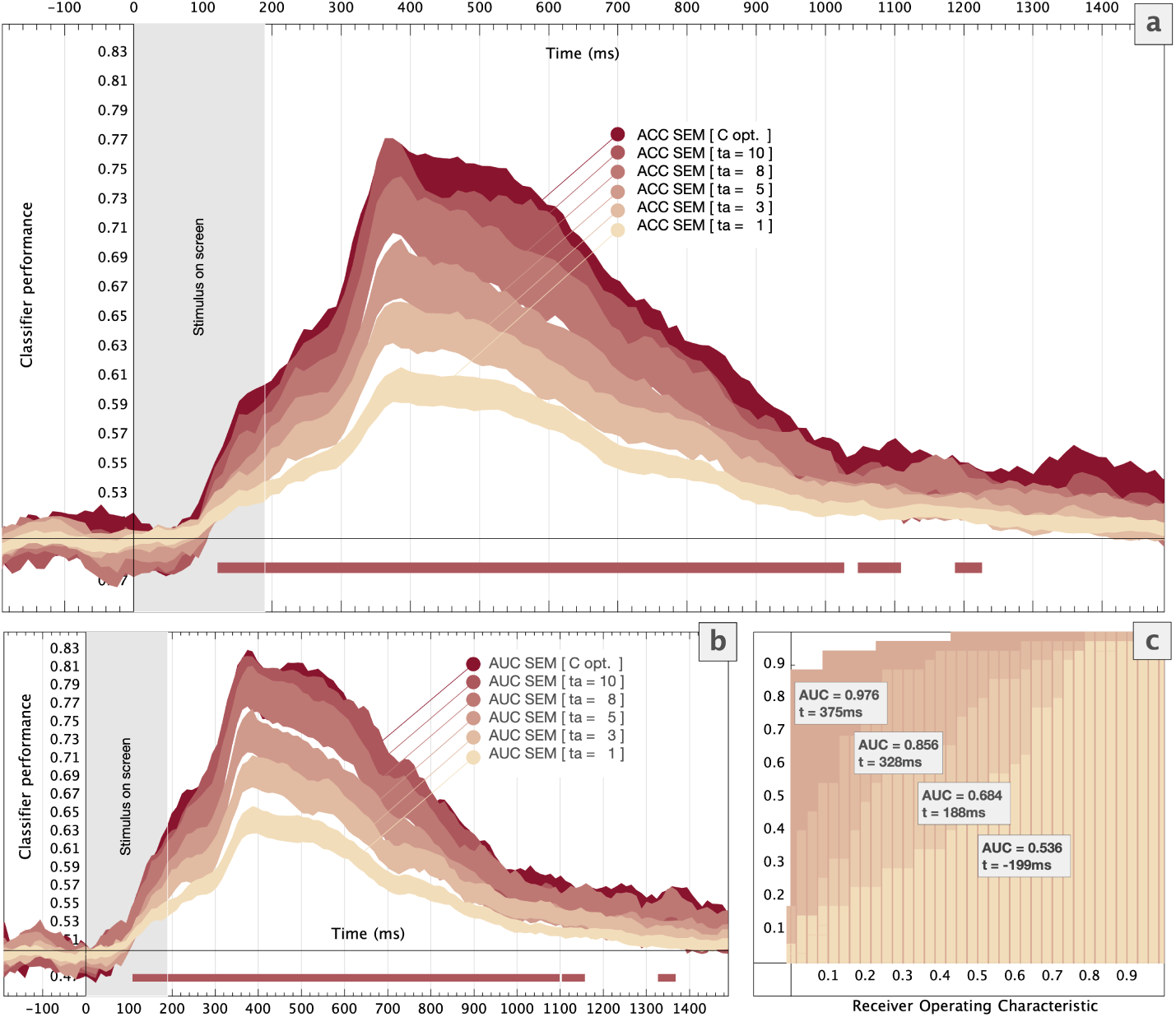
Group level MVPA results. Time-resolved classifier performance when different number of trials were averaged. The standard error of the **(a)** classification accuracy and **(b)** the Area Under the Curve are represented using colored areas. Significant windows (*t*_*a*_ = 10) obtained via Stelzer permutation test are highlighted using horizontal bold lines. The stimulus screen time [0 − 190]ms is shaded. **(c)** Receiver Operating Characteristic curves for different timepoints [example participant, *t*_*a*_ = 10, *t* =-199ms, 188ms, 328ms and 375ms].

The statistically significant regions extended from 130ms after stimulus onset to 1200ms afterwards, when ten trials were averaged to generate supertrials. As Figure 5 shows, before the stimulus onset the classification accuracy remained at the chance level (0.5).

The temporal generalization analysis is shown in Figure 6. First, AUC proved to be a more sensitive measure. The AUC temporal generalization matrix (Fig 6b) shows a distinct pattern of generalization. Clusters appearing only alongside the diagonal have been associated with a succession of different mechanisms. That is to say, the neural information that allows the classifier to tell apart congruent and incongruent trials is likely the result of a series of distinct events. Moreover, Figure 6b shows a cluster of homogeneous AUC between 200 and 400ms, which theoretically suggests the operation of a single cognitive process maintained in time.^34^ Such mechanism apparently reappears at ∼ 800-1000ms after the target onset, posterior to the mean RT (513ms). In consequence, results indicate the existence of a particular brain process involved in the interference effect that intervenes in the initial stages of target processing during an extended time window, and reappears after the behavioral response is given.

**Figure 6:**
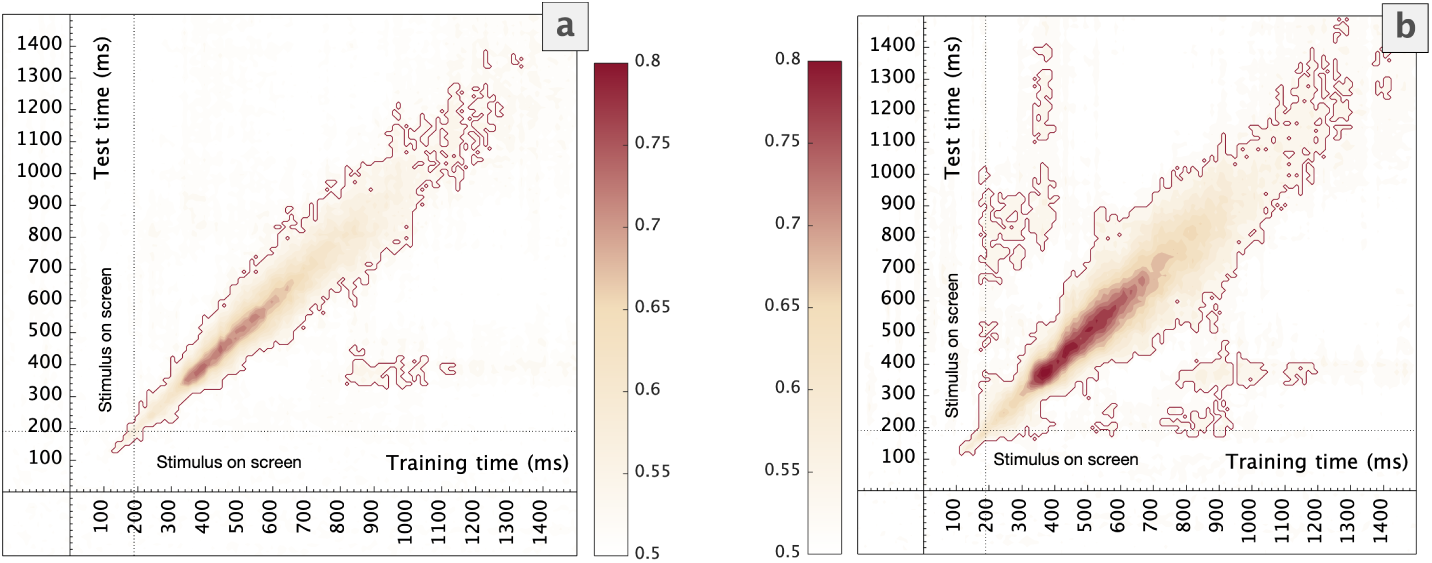
Group level temporal generalization results for congruent vs. incongruent trials (*t*_*a*_ = 10). Accuracy **(a)** and AUC **(b)** values when the model was trained and tested in each time point of the whole time window. Significant clusters obtained via Stelzer permutation tests are highlighted using red lines.

**Figure 7:**
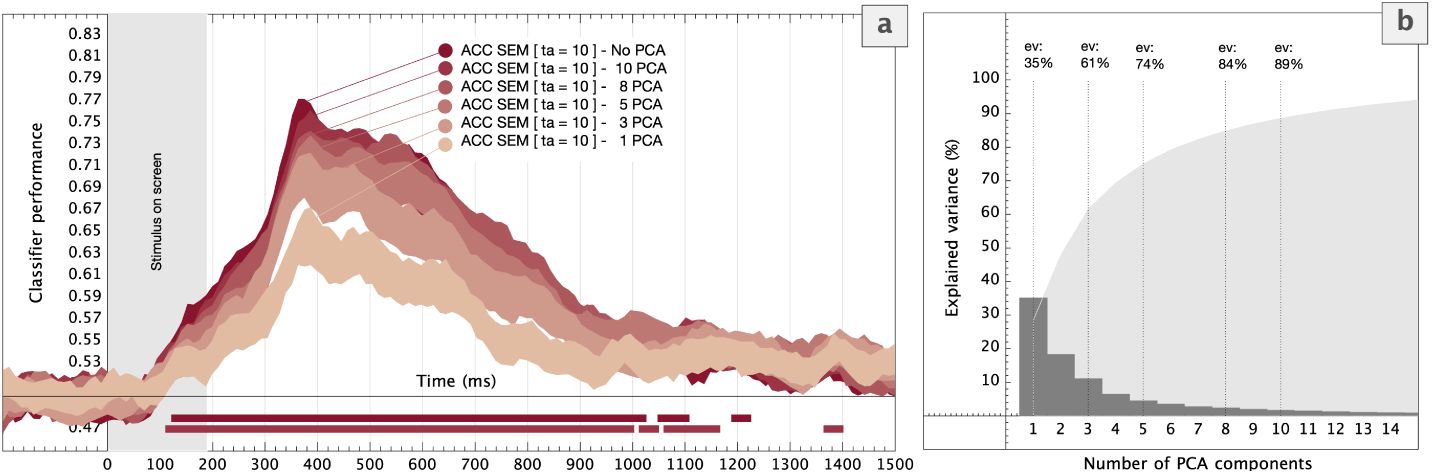
Group level MVPA. **(a)** Time-resolved classifier performance (ACC) for congruent vs. incongruent trials when different number of PCA components were selected. Colored areas represent the ACC standard error. Statistically significant time windows (*t*_*a*_ = 10, No PCA and 10 PCA comp.) are highlighted using horizontal bold lines. The stimulus screen time [0-190]ms is shaded. **(b)** Explained variance for different number of PCA components [example participant].

**Table 2:** LSVM model peak classification performance [*t* = 375*ms*] at a group level. The mean accuracy and AUC are reported for different values of *t*_*a*_ and different number of PCA components.

| No. of averaged trials ($t_a$) | ACC ± SD | AUC ± SD |
| --- | --- | --- |
| $t_a = 1$ | .60±.05 | .65±.07 |
| $t_a = 3$ | .65±.07 | .70±.08 |
| $t_a = 5$ | .69±.10 | .74±.12 |
| $t_a = 8$ | .74±.10 | .79±.12 |
| $t_a = 10$ | .76±.11 | .80±.13 |
| $t_a = 10$ - C optimized | .76±.10 | .81±.13 |

| No. of PCA components ($t_a = 10$) | | |
| --- | --- | --- |
| First Component | .64±.14 | .66±.17 |
| 3 first components | .71±.11 | .76±.13 |
| 5 first components | .72±.11 | .78±.12 |
| 8 first components | .73±.12 | .80±.13 |
| 10 first components | .74±.11 | .81±.13 |

Interestingly, this is the same temporal window where classic Event-Related Potential studies^52^ have repeatedly observed the N2 potential, which is taken as the reflection of interference processing. The indication that the same underlying mechanism reappears after the response could reflect the reinstantiation of the interference episode, perhaps reflecting trial event boundaries.^53^ Further research will be needed to clarify and extend this novel finding.

The cross-classification results (Figure 9a,b) showed smaller clusters compared to the MVPA time generalization (Figure6a, b). However, the main diagonal cluster in the matrix indicates a series of different events that occur in cascade, but shared between both contexts.^34^ This mechanism could reflect the interference process itself, previous to the response.

The actual risk estimation for different sample sizes and dimensions *γ*_*emp*_ is shown in Figure 8. The difference between the actual error and the resubstitution error is bounded by *γ*_*emp*_. White markers represent different experimental configurations for both the sample size (*l*) and the number of PCA components (*d*) analyzed in our study. Performance results obtained by resubstitution (C-optimized, *t* = 375ms) for these experimental configurations are shown in Table 3. The classification accuracy remained above chance despite the conservative estimation of the upper bound of the actual error, preserving our classification model for overfitting and proving that both conditions are representative of different underlying activity patterns associated with congruent and incongruent stimuli.

**Table 3:** Classification performance and the actual risk *γ*_*emp*_ for a C-optimized LSVM model obtained by the resubstitution approach. [example participant, *t* = 375ms]

| $\text{ACC}(\gamma_{emp}(l, d))$ | $d = 1\text{PCA}$ | $d = 3\text{PCA}$ | $d = 5\text{PCA}$ |
| --- | --- | --- | --- |
| $l = 790$ ($ta = 1$) | .55(.04) | .63(.10) | .65(.13) |
| $l = 260$ ($ta = 3$) | .58(.08) | .68(.16) | .71(.20) |
| $l = 150$ ($ta = 5$) | .58(.11) | .79(.20) | .81(.25) |
| $l = 90$ ($ta = 8$) | .72(.14) | .80(.24) | .83(.30) |
| $l = 70$ ($ta = 10$) | .81(.15) | .88(.27) | .92(.34) |

**Figure 8:**
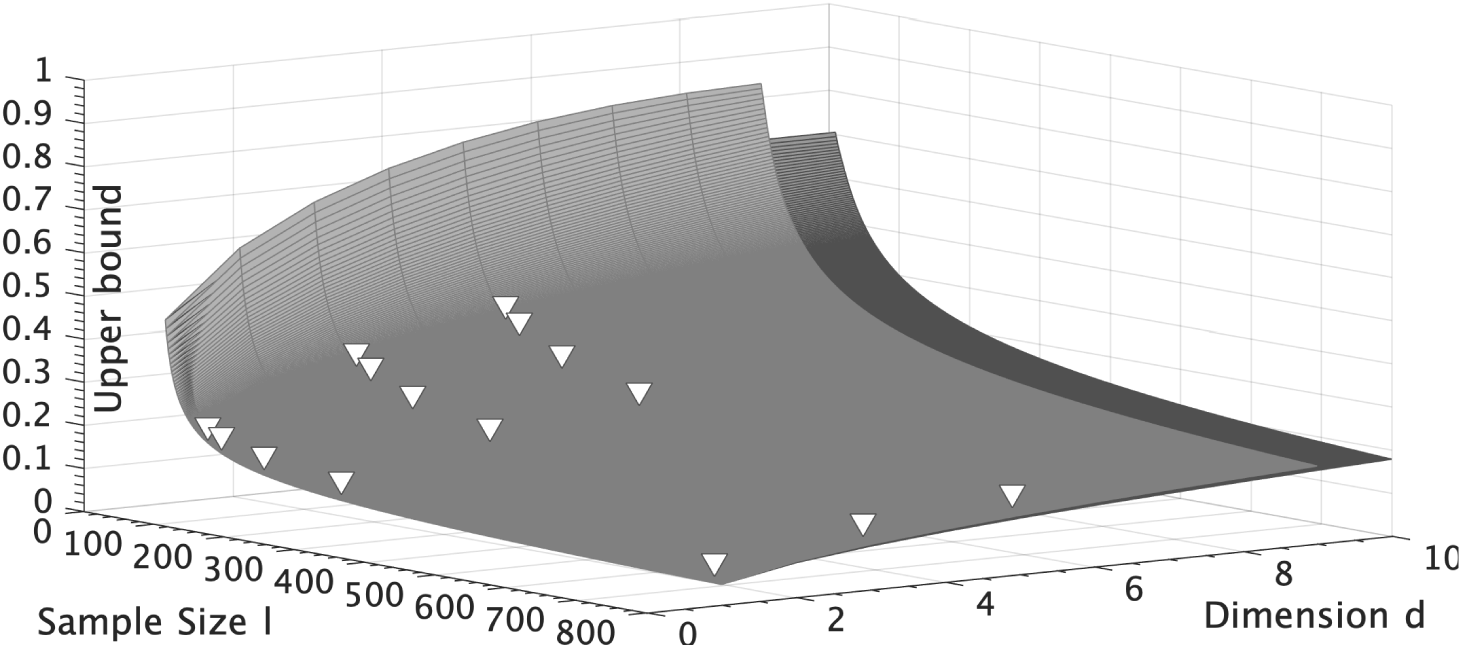
Different upper bound estimations via the procedure found in^35^ for LSVM across dimension and sample size at a 95% confidence level (*η* = 0.05). White markers represent the upper bound values for the experimental conditions tested in our study.

**Figure 9:**
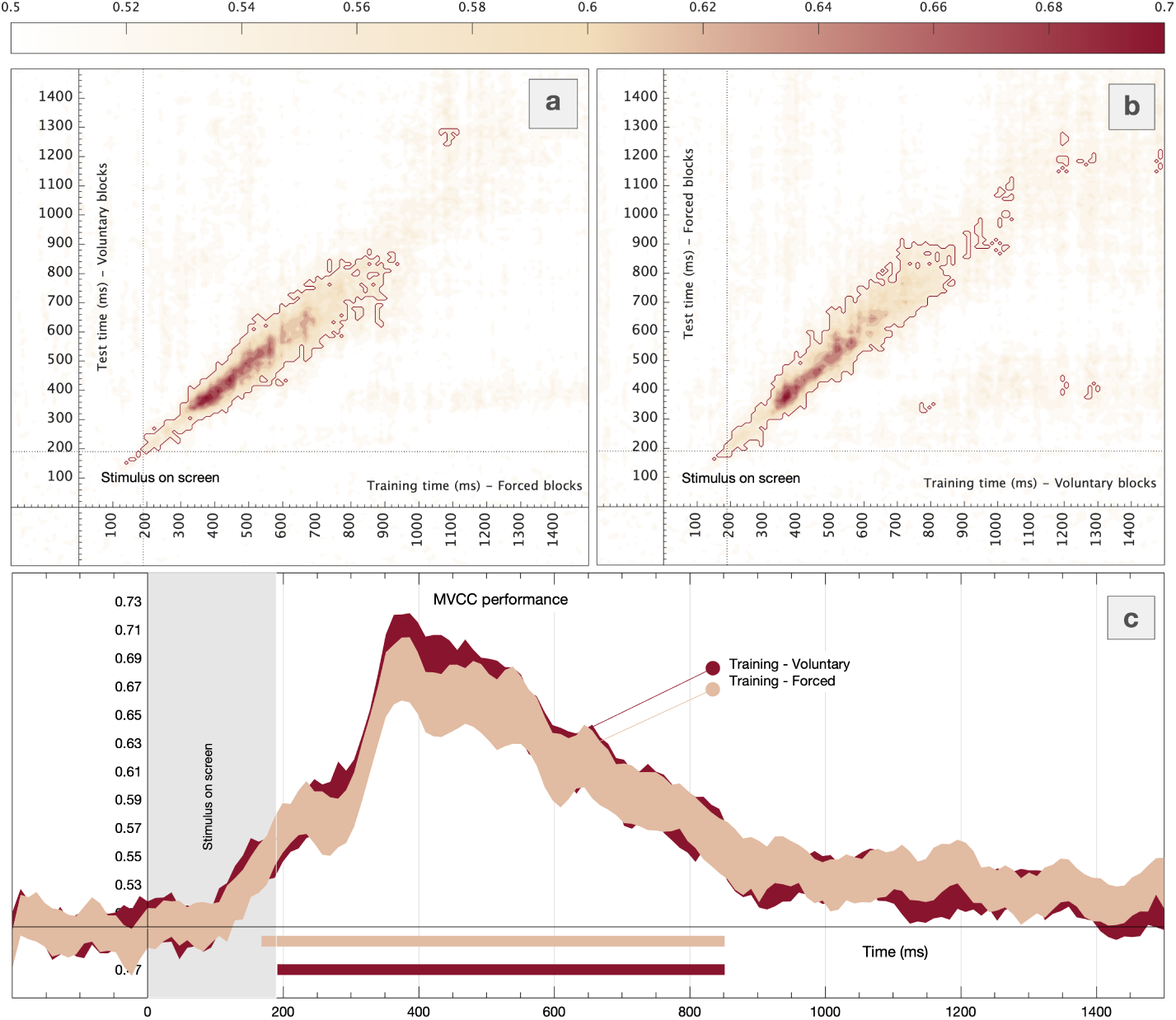
MVCC results. **(a)** Temporal generalization results when the model was trained with forced blocks and tested in voluntary blocks and **(b)** *vice versa*. **(c)** Classifier performance (acc) for the former analyses. Colored areas represent the standard error. Significant windows calculated via Stelzer permutation test are highlighted using horizontal lines.

### 3.3 Frequency contribution results

A sliding bandstop filter approach was followed to study the contribution of each frequency band to the overall decoding accuracy. Results show that the interference effect observed relies on neural processes operating in the Delta and Theta frequency bands. Figure 10a shows how decoding accuracy significantly drops when frequencies up to 8Hz were filtered-out.

Previous studies on cognitive control, and more specifically, interference processing, have found that slow rhythms (i.e. Theta, Delta) are associated with communication between distant brain regions.^54^ Our results are in line with those studies, showing that Theta and Delta oscillations are relevant for the brain activity underlying performance in an interference task. Moreover, previous results (e.g. Cohen^55^) show the relevance of Theta in the first instances of target processing, which changed to Delta after the participants’ response. These results are supported by the present study, which shows Theta and Delta to be crucial for classification right after the target onset, which evolves into a single Delta-based classification around and after the response time.

The meaning of the change from one frequency band to another along time could be due to neuronal activity on the Theta band preventing the distractors to be processed. Once the target is selected, Delta, which arises later, could reflect inhibition of competing and erroneous motor responses.^56^

## 4 Conclusion

The current study is an initial approximation to adapt a DST to a format that allows measuring concurrent high-density electroencephalography. While most of previous studies categorize the interference effect through ERP markers such as the N2 potential,^52^ we successfully used multivariate pattern analysis (MVPA) to decode conflict-related neural processes associated with congruent or incongruent events in a time-frequency resolved way. Furthermore, our results of frequency bands contribution analysis suggest that interference processing effect relies on neural processes operating in the Delta and Theta frequency band. This is in line with previous results pointing to Theta band modulations as a cognitive control signature, indicating the presence of conflicting/incongruent information. More specifically, the source of this modulation has been localized in the midline brain, in anterior cingulate and premotor areas.^53^

In addition, our results offer new information regarding the reinstantiation of the processes likely reflected on the N2 potential at a later time window in the trial. This finding, which could not have been obtained with classic analytical strategies, opens novel avenues of research. Future lines of investigation should address these findings to complement the results found in the current investigation. In addition, to increase our understanding of preparation processes and conflict effects, it would be of interest to continue analyzing the current dataset, focusing not only on the target stimulus, but also on the neural activity triggered by the cues. Further detailed analyses should be carried out to study the activation differences between forced and voluntary blocks or high and low congruency contexts. MVPA techniques represent an opportunity to study the neural basis of even more complex psychological processes, such as the allocation of cognitive control and effort avoidance. Finally, the use of the resubstitution approach mentioned above opens a new path that could lead to promising results.

**Figure 10:**
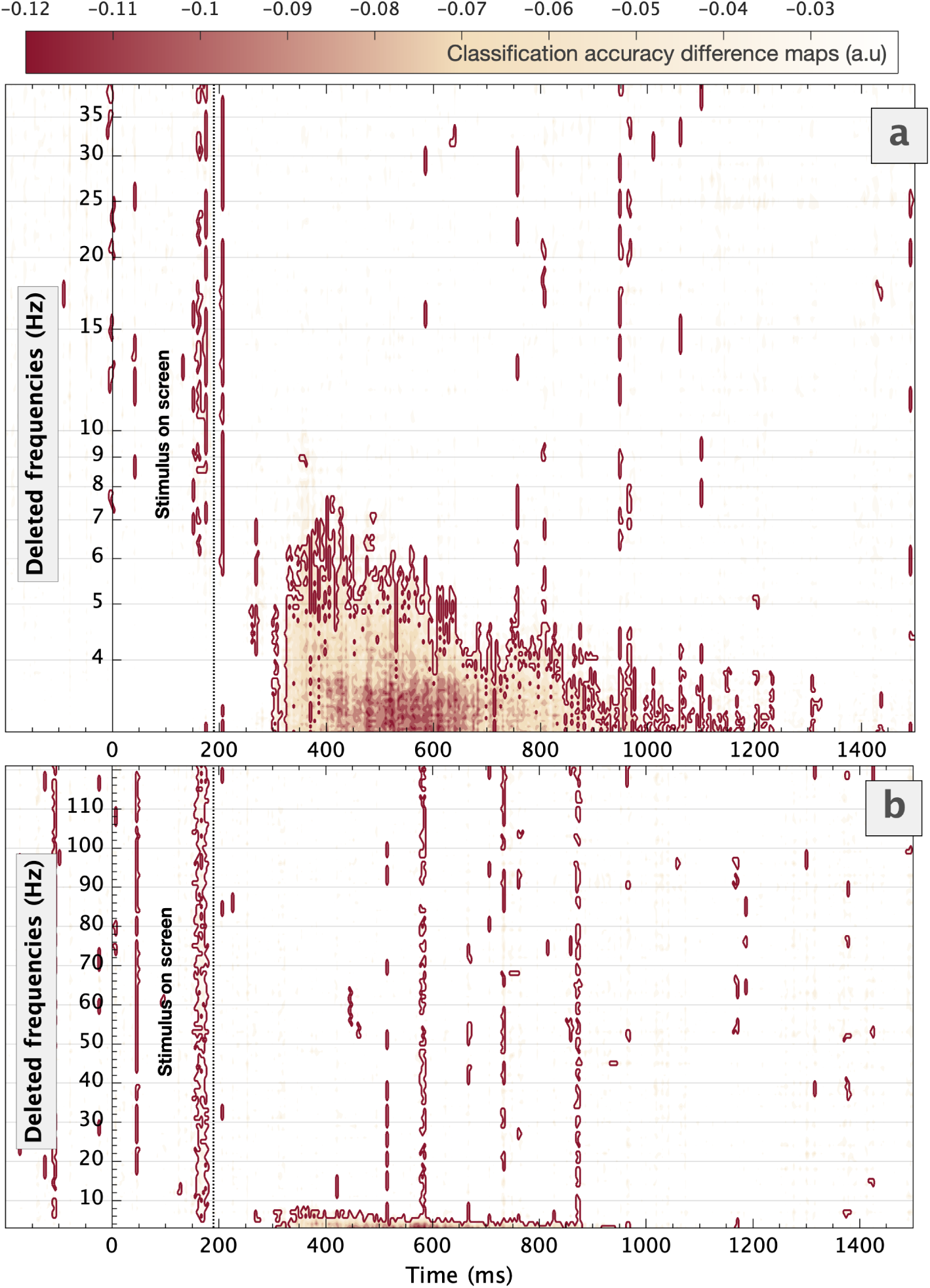
Results of frequency contribution analysis. Classification accuracy differences when a specific frequency band is filtered-out. **(a)** [0-40]Hz logarithmically spaced and **(b)** [0-120]Hz linearly spaced sliding filter approach. Significant clusters obtained via Stelzer permutation test are highlighted using red lines.

## Acknowledgments

This research was supported by the Spanish Ministry of Economy and Business under the TEC2015-64718-R and PSI2016-78236-P grants. The first author of this work is supported by a scholarship from the Spanish Ministry of Economy and Business (BES-2017-079769).

